# Seeing clearly with CLARI-O: a window into cellular architecture, interactions, and morphology of organoid models

**DOI:** 10.64898/2026.03.29.715075

**Authors:** Samra Beyene, Martin Thunemann, Elizabeth K. Kharitonova, Natalie Baker Campbell, Shira Klorfeld-Auslender, Farzad Mortazavi, Ella Zeldich

**Author notes:** co-senior authors (Farzad Mortazavi, Ella Zeldich).

## Abstract

Cortical organoids (COs) represent a powerful in vitro model system that recapitulates key aspects of human brain development, enabling the study of neurodevelopmental processes, cellular diversity, and disease mechanisms in a physiologically relevant 3D environment. However, traditional histological analysis of COs relies on tissue sectioning, which limits the ability to capture the full spatial complexity of organoid architecture. In this study, we establish a framework for applying CLARI-O, an improved tissue-clearing technique, for intact COs and organoid-based systems, enabling comprehensive 3D visualization and analysis of 3D organizational features. Using CLARI-O in combination with high-resolution imaging, we demonstrate the utility of tissue clearing for studying glial populations, including oligodendrocytes and microglia, considered to be underrepresented in COs, and their interactions with neurons. Additionally, we apply this method to forebrain assembloids (FAs) to visualize cellular heterogeneity and the interface between ventral and dorsal regions. Finally, we use CLARI-O to study mouse brains containing xenotransplanted COs (MB-COs) to evaluate human cell integration, migration, vascularization, and structural connectivity. This is the first study to demonstrate how tissue clearing can be used after functional assays such as calcium imaging to correlate neural activity with post hoc structural analysis in MB-COs. Together, this work establishes CLARI-O as a powerful tool for advancing 3D structural and functional interrogation of human CO-derived systems, enhancing their value for disease modeling, drug screening, and translational neuroscience.

**Motivation:** Cortical organoids have become an increasingly powerful tool in neuroscience. Their complexity has expanded substantially, now incorporating exogenous lineages, fusing organoids with distinct regional identities (assembloids), and enabling xenotransplantation into *in-vivo* environments. These advancements require more sophisticated technological approaches that are capable of capturing the intricate three-dimensional cyotarchitecture and organization of intact organoid systems both in vitro and after xenotransplantation *in vivo*. Tissue-clearing methodologies offer a unique opportunity to visualize these structural and cellular features with exceptional depth and resolution.

**Graphical abstract:** 

**Highlights:**

- We optimized clearing protocols to develop an organoid specific clearing method (CLARI-O) that enables high-resolution visualization of diverse neuronal and glial populations without tissue sectioning, preserving long-range connections and cellular processes.
- Forebrain assembloids used to study neuronal and oligodendrocyte migration can be effectively processed using CLARI-O, allowing detailed visualization of fusion interface.
- We established a robust framework for CLARI-O-based clearing of mouse brain tissue containing xenotransplanted human cortical organoids, enabling comprehensive 3D analysis of graft development, integration, and vascularization in vivo.

## Introduction

The three-dimensional (3D) nature of human cortical organoids (COs) uniquely offers a holistic view of developmental pathways, cellular kinetics and interactions, connectivity, and functional dynamics in a single model. COs are generated through self-assembly of either embryonic stem cells or induced pluripotent stem cells (iPSCs), including patient-derived iPSCs. These COs can be regionally patterned to represent more region-specific cell types, which in turn form organized structures composed of progenitor, neuronal, and glial cell types. Such cells transcriptionally and electrophysiologically resemble mid-term gestation human prenatal brains^1–8^. Within these 3D platforms, multiple cell lineages differentiate, migrate, mature, and interact with each other. This allows for researchers to study the intricate dynamics of tissue organization, development, and function in ways that are not possible in traditional 2D cultures. Recent advances have further extended the utility of COs by enabling their xenotransplantation into rodent brains, where human cells achieve unprecedented levels of maturation and functional integration into host cortical circuits^9,10^.

A widely used method for characterizing organoid morphology and cell composition is immunohistochemical analysis of fixed serial sections^5,11–13^. This traditional histological approach relies on thin slicing of tissue, which offers only a limited, two-dimensional view, thus, offering a limited representation of the organoid-wide neural networks, structural connectivity, and complex internal organization. In addition, mechanical sectioning inevitably severs axonal projections and long-range processes, obscuring the true three-dimensional architecture of neural circuits. Furthermore, the effectiveness of immunohistochemical analysis can be compromised by the low permeability of antibodies and potential tissue damage during sectioning. These limitations restrict our ability to take full advantage of the complex cytoarchitecture and spatial arrangements within COs in vitro and mouse brains containing xenotransplanted COs (MB-COs) *ex-vivo*. To overcome these limitations, a sectioning-free, whole-organoid imaging platform is needed.

Tissue clearing techniques have significantly advanced organoid research by allowing scientists to visualize the intricate 3D structures within these miniaturized organ models^14,15^. By removing lipids that cause light scattering, tissue clearing makes entire organoids transparent, while preserving proteins and overall structural integrity^16^, thus enabling complete, three-dimensional imaging and reconstruction without the need for physical sectioning. This allows for the visualization of immunofluorescently labeled membrane, synaptic, signaling, and lineage-specific marker proteins. We had previously published a detailed clearing protocol in 2019 ^17^, which we validated in zebrafish, mouse, rat, non-human primate, and human brain samples. Here, we describe how we have optimized this protocol and developed CLARI-O, an organoid-specific clearing protocol that maintains structural integrity, preserves cell-specific fluorescent labeling, and enables repeated imaging of the same sample. Specifically, we adapted the 2019 protocol^17^ by incorporating low-speed centrifugation and extended heat-assisted passive clearing into the workflow, which improved clearing uniformity across samples of varying size and complexity while minimizing mechanical disruption. Furthermore, these refinements improved reagent penetration while preserving tissue integrity and endogenous fluorescence, ensuring high-fidelity imaging across multiple rounds of imaging and analysis.

CLARI-O was used to clear COs, forebrain assembloids (FAs), and MB-COs, enabling high-resolution, three-dimensional analysis of organoid architecture and the longitudinal integration of CO-derived human cells within the mouse brain following *in-vivo* functional assays. Using CLARI-O, we were able to visualize not only the canonical cell populations present in dorsally derived organoids, such as neurons and astrocytes, but also cell types not typically represented in classical CO models. Here, we demonstrated that our CLARI-O approach allows for the visualization of distinct niches of oligodendrocytes that emerge following expansion of glial progenitors, and exogenously incorporated iPSC-derived microglia, as well as the detailed interactions between neuron and glia cells. Because these interactions are spatially organized, preserving intact three-dimensional structure is essential for accurate interpretation.

Furthermore, xenotransplantation of COs into the mouse cortex provides a maturation-promoting environment that supports vascularization and enables a more comprehensive assessment of neuronal activity. Following implantation of virally labeled cells and in vivo calcium imaging, the host mouse brains were subsequently cleared to allow volumetric analysis of the graft and surrounding tissue. CLARI-O–based imaging enabled visualization of the overall MB-CO architecture, its vascular integration, and the spatial distribution of human cells within the host brain. Together, the integration of functional live-cell imaging with high-resolution 3D structural analysis through CLARI-O provides a powerful and complementary strategy to dissect structure–function relationships, elucidate disease mechanisms, evaluate therapeutic interventions, and advance next-generation models of human brain development.

Altogether, this study establishes a robust and scalable framework for leveraging CLARI-O to comprehensively visualize and interrogate the structural organization, cellular diversity, and integrative complexity of COs and MB-COs within their intact three-dimensional environment.

## STAR Methods

### Key resources table

#### iPSC lines and maintenance

We used a female iPSC line with a normal karyotype (WC-24-02-DS-B) utilized in our previous work^13,18^, a male control cell line (BU3-10-Cr2) that was obtained from the Center of Regenerative Medicine (CReM), Boston University School of Medicine, and a female cell line MGH2046^19^ kindly provided by Dr. Steven J. Haggarty (Harvard Medical School and Massachusetts General Hospital). The cells were cultured on Matrigel (cat. 354277 Corning^®^, Corning, NY, USA) and passaged using ReLeSR (cat. 85850, STEMCELL Technologies, Vancouver, BC, Canada). The mTeSR plus (cat. 85850, STEMCELL Technologies) media was changed every other day. iPSCs between passages 24 to 50 with normal morphology were used for the organoid generation.

### Generation of cortical organoids and assembloids

#### Generation of dorsally derived COs (dCOs)

COs were generated following the protocol established by Pasca’s group^5^ with small modifications. Briefly, one well of iPSCs from a 6-well plate was washed with 1 mL of PBS. The dissociation of cells was achieved via treatment with 1 mL of Accutase (cat. 7920, STEMCELL Technologies) for ∼5 min. The resuspended dissociated single cell suspension was combined with 5 mL of DMEM/F12 (cat. 11330–057, ThermoFisher Scientific, Waltham, MA, USA), centrifuged for 5 min and resuspended in 1 mL mTeSR plus with 50 µM of Rock Inhibitor (cat. 12-541-0, Fisher Scientific, Pittsburgh, PA, USA) for counting. The cells were plated in low adherence V-bottom 96 well plates (cat. MS-9096VZ, S-Bio Prime, Constantine, MI, USA), at a density of 15,000 cells/well/150 µL of media. The next day (day 1), the media was replaced with 100 µL of TeSR-E6 (cat. 0596, STEMCELL Technologies) that was supplemented with 2.5 µM of Dorsomorphin (cat. P5499, MilliporeSigma, Burlington, MA, USA) and 10 µM SB-431542 (cat. S4317, MilliporeSigma). 100 µL of TeSR-E6 media containing aforementioned concentrations of Dorsomorphin and SB-431542 was replaced daily between days 3 – 6.

On day 7, 100 µL of the media was switched to Neural Media (NM) containing B-27 supplement, minus vitamin A (1:50; cat. 12587, Life Technologies), GlutaMax (1:100, cat. 35050061, Life Technologies), 100 U/mL Penicillin/Streptomycin (1:100, cat. 15140122, Life Technologies) and Primocin (1:500; cat. ant-pm-1, Invivogen, San Diego, CA, USA) in Neurobasal A Medium (cat. 10888022, Life Technologies, Carlsbad, CA, USA). The NM was supplemented with fibroblast growth factor 2 (FGF2; 20 ng/mL, cat. 233-FB-25/CF, R&D Systems, Minneapolis, MN, USA) and epidermal growth factor (EGF; 20 ng/mL, cat. 236-EG-200, R&D Systems) between days 7 – 25. COs were transferred to 24-well ultra-low attachment plates (cat. 07-200-602, ThermoFisher Scientific) between days 16 to 24 and cultured individually. From days 25 to day 39, NM with brain-derived neurotrophic factor (BDNF; 20 ng/mL, cat. AF-450-02, PeproTech, Waltham, MA, USA), neurotrophic factor 3 (NT3; 20 ng/mL, cat. 450-03, PeproTech) and 1% Geltrex (cat. A1569601, Life Technologies) was used. Half media changes were performed every other day. From day 40 onwards, dCOs were maintained in NM with 1% Geltrex, with a half change media performed every two days.

#### Generation of dCOs enriched with oligodendrocytes (dOCOs)

dOCOs were generated as we described before^13,20^. The dCO protocol was followed until day 50. To expand the oligodendrocyte population within dCOs, NM was supplemented with platelet-derived growth factor AA (PDGF-AA; 10 ng/mL, cat. 221-AA, R&D Systems) and insulin-like growth factor (IGF; 10 ng/mL, cat. USA291-GF-200, R&D Systems) between days 50-60. Between days 61 to 69, 3,3′,5-triiodothronine (T3; 40 ng/mL, cat. T6397, MilliporeSigma) was added to the NM and the media was changed every two days. From day 70 onwards, NM was changed every two days.

#### Generation of ventrally derived COs (vCOs)

The generation of vCOs was achieved through ventral patterning factors and modifications to the dCO protocol. The generation of the vCOs followed the same steps as dCOs between days 0-6 except for the Wnt pathway inhibitor IWP-2 (5 μM, cat. S7085 Selleckchem, Houston, TX, USA) that was added between days 4 and 24. Starting on day 7, vCOs were transitioned to the differentiation and maintenance medium (DMM)^21^ containing DMEM/F12, B-27 supplement, minus vitamin A (1:50), N2 (cat. 17502048, ThermoFisher Scientific), human insulin (25 μg/ml, cat. I9278-5ML, MilliporeSigma), non-essential amino acids (NEAA; 1:100, cat. 11140076, ThermoFisher Scientific), Penicillin/Streptomycin (1:100), GlutaMax (1:100), and β-mercaptoethanol (0.1 mM, cat. M3148, MilliporeSigma). EGF (20 ng/ml) and FGF2 (20 ng/ml) were added to DMM media between days 7 to 24. The Sonic Hedgehog pathway agonist, smoothened agonist (SAG; 1 μM, cat. 566660, MilliporeSigma) was supplemented to the DMM between days 12 to day 24. The neuronal expansion and enrichment for oligodendrocytes within vCOs was achieved through the supplementation of DMM media with BDNF (20 ng/mL), NT3 (20 ng/mL), PDGF-AA (10 ng/mL), IGF (10 ng/mL), T3 (60 ng/mL), Hepatocyte growth factor (HGF; 5 ng/mL, cat. 315–23, PeproTech), cyclic AMP (cAMP; 1 μM, cat. D0627, MilliporeSigma), and biotin (100 ng/mL, cat. B4639, MilliporeSigma) between days 24 to 37^21^. Starting from day 37, vCOs were cultured in DMM supplemented with T3, cAMP, biotin, and ascorbic acid (AA; 20 μg/mL, cat. A4403, MilliporeSigma), and media was changed twice a week.

#### Generation of Forebrain Assembloids (FAs)

To generate FAs, on day 70 vCOs and either dCOs or dOCOs were transferred to microcentrifuge tubes with 1 mL of NM media and kept in an incubator for 3 days. A half-volume media change was performed gently on the second day. Following the fusion, FAs were transferred into 24-well ultra-low attachment plates, and the NM was half-changed every other day^22^.

#### Lenti virus (LV) labeling of dCOs

LV labeling of COs for imaging following xenotransplantation was done similarly to AAV labeling. pLV-Puro-SYN1.jGCaMP8s (cat. VB240904-1449tnm, VectorBuilder, Chicago, IL, USA) and pLV-Bls-EF1A>mScarlet3 (Cat. VB240919-1466xpw, VectorBuilder) were used at a titer of ≥ 1×10^9^ TU/mL. Starting on 40 DIV, COs were moved to a 1.5-mL tube and incubated for 30 minutes at 37 °C with 2 μL of pLV-Bls-EF1A>mScarlet3 and 4 μL of pLV-Puro-SYN1.jGCaMP8s and ∼20μL of maintenance media. Following overnight incubation in 300 μL of maintenance media, COs were moved back to 24-wells for maintenance until grafting procedure. Starting on 40 DIV, verified fluorescent COs were used for transplantations.

#### Microglia generation

iPSCs were differentiated into hematopoietic progenitor cells (HPCs) for 12 days using the STEMdiff Hematopoietic Kit (Catalog #05310, STEMCELL Technologies) by following the manufactural instructions. On day 12, hematopoietic cells (HPCs) were collected and the expression of CD43, CD45, and CD34 was validated by flow cytometry. Following this, 700,000 HPCs were cryoprotected in 1mL of freezing media (cat. 05859, STEMCELL Technologies). For the generation of microglia, 200,000 HPCs were thawed and for 24 days they were differentiated into microglia using the STEMdiff Microglia Differentiation Kit and culture protocol (cat. 100-0019, STEMCELL Technologies) per manufactural instructions. On day 24, the expression of CD45, CD11b, and TREM2 was validated by flow cytometry and the co-expression of CD45 and CD11b in more than 80% of the cells were considered a successful differentiation.

#### Microglia incorporation into COs

To incorporate exogenous microglia into the corresponding COs, 24-day microglial progenitors were allowed to spontaneously migrate into 50-day-old COs. For the incorporation, a single CO was transferred to an eppendorf tube with the carryover medium from the CO. After this, 50,000 microglia cells from the same line were suspended in 100 μL of microglia differentiation medium from the STEMdiff Microglia Differentiation Kit and culture protocol (cat. 100-0019, STEMCELL Technologies) added to the eppendorf tube. COs were incubated with the microglia in the Eppendorf tubes for 3 days to allow the microglia to attach to the surface of the COs. After this 3-day incubation, the COs with the attached microglia were transferred back to the ultra-low adherence 24-well plates. They were cultured until 8 days after microglia incorporation in the Neural Media with 1% Geltrex with the addition of STEMdiff Microglia Supplement 2 (cat. 100-0023, StemCell Technologies) at a 1:250 dilution^23,24^. On day 8 following incorporation the microglia containing COs were fixed in 4% Paraformaldehyde (PFA, cat. P6148, MilliporeSigma) as described below.

#### Fixation

Different types of COs and FAs were transferred to Eppendorf tubes, washed once with PBS, and then transferred to 15 mL Falcon tubes with approximately 5 mL of 4% PFA for 48 hours of fixation.

#### Xenotransplantation and in-vivo imaging

Experiments and animal procedures were conducted according to the Guide for the Care and Use of Laboratory Animals and protocols approved by the Boston University Institutional Animal Care and Use Committee (PROTO202000026). COs were transplanted into the mouse retrosplenial cortex as we described previously^9^. Briefly, we used non-obese diabetic, severe combined immunodeficient (NOD/SCID) mice (Charles River Laboratories) at an age of 12-20 weeks that were single- or group-housed in isolated ventilated rodent cages under normal light-dark cycles with nestlet enrichment and unrestricted access to food and water. Animals received 4.8 mg/kg Dexamethasone, 1 mg/kg extended-release Buprenorphine, and 5 mg/kg Meloxicam 120-180 min before surgery. Animals were anesthetized with a cocktail of 0.05 mg/kg Fentanyl, 0.25 mg/kg Dexmedetomidine, and 5 mg/kg Midazolam, were placed within a stereotaxic frame on a feedback-controlled heating blanket and received 100% oxygen through a nose cone for the duration of the surgery. After skin removal and cleaning of the dorsal cranium, we placed a titanium headbar, performed craniotomy (3-mm diameter) and durectomy, aspirated a ca. ∼1 mm^3 large piece of retrosplenial cortex, placed a single hCO in the resulting cavity, and closed the exposure with a glass window. Immediately after surgery, animals received a single dose of 500 mg/kg Cefazolin; Meloxicam (5 mg/kg every 24 h) was administered for three days after surgery. Animals received Sulfamethoxazole/Trimethoprim (Sulfatrim) via medicated food (Uniprim diet, Envigo, TD06596) for the duration of the study.

After a recovery period of seven days, animals underwent habituation in one session per day to accept increasingly longer periods (up-to 120 min) of head restraint inside the microscope enclosure. Drops of sugar water (infusion-grade 5% dextrose in 0.9% NaCl) were offered as a reward during training and recording sessions. Two-photon imaging was performed in awake, head-fixed animals on a commercial two-photon laser scanning microscope system (Bruker Ultima Investigator Plus) with Coherent Chameleon Discovery Ti:Sapphire lasers tuned to 920-950 nm for excitation. For vascular imaging, in-house produced Alexa 680-Dextran (using amino-dextran with 0.5 MDa molecular weight, Fina Biosolutions) was injected under isoflurane anesthesia 15-30 min before imaging. Overview images of the cranial window were acquired with a 4× objective (Plan-Neofluar, NA = 0.16, Zeiss) at the beginning of each imaging session. For functional imaging, a 20× objective (XLUMPlanFLNXW, NA = 1.0, Olympus) was used. Calcium imaging was performed in consecutive 180-300-s long acquisition runs in galvo/resonant galvo mode with an image size of 512 × 512 pixels and an acquisition rate of 15-30 Hz. Image time series underwent in-plane motion correction with ‘normcorre’ (Pnevmatikakis and Giovannucci 2017) using the reference channel showing mScarlet fluorescence; then, Suite2P (Pachitariu et al. 2017) was used for automatic ROI identification with Cellpose using the setting ‘anatomical_only=2’ and extraction of fluorescence changes in neuronal cell bodies as ΔF/F.

### CLARI-O Procedure

#### For COs and FAs

following fixation, COs and FAs were immediately transferred to clearing solution consisting of 200 mM sodium dodecyl sulfate (SDS; cat. 1610302, Bio-Rad, Hercules, CA, USA) and 20 mM lithium hydroxide monohydrate (cat. L127500, Fisher Scientific, Pittsburgh, PA, USA) dissolved in 1 M boric acid (cat. B2645, Sigma-Aldrich, Burlington, MA, USA). The pH of the clearing solution was adjusted to 9.0 using 0.1 M sodium hydroxide. Samples were placed in 15-mL Falcon tubes and incubated in a 37 °C water bath for 10 days to promote effective lipid extraction.

During the 10-day clearing period, the clearing solution was replaced approximately every 3 days with freshly prepared clearing solution that had been pre-warmed to 37 °C. Maintaining the clearing solution at physiological temperature is critical; transferring the samples into cold or room-temperature clearing solution substantially slows lipid solubilization and can extend the clearing timeline.

On day 10, COs and FAs were centrifuged in pre-warmed clearing solution for 5 minutes at 1500 RPM at 30 °C, followed by the incubation in the fresh clearing solution at 37 °C for additional 5 days to achieve optical transparency, with the clearing solution replaced every 3 days with freshly prepared clearing solution that had been pre-warmed to 37 °C. On day 15, samples were transferred to a fresh, pre-warmed clearing solution and centrifuged for 5 minutes at 1500 RPM at 30 °C. Then, the samples were transferred to directly to 0.05 M Tris-Buffered Saline (TBS), pH 7.6 (cat. PPB011, Sigma-Aldrich, Burlington, MA, USA) pre-warmed to 37 °C and brought to room temperature.

After transfer, COs and FAs were kept on a shaker in 0.05 M TBS at room temperature until immunostaining for up to 2 months. The timeline described here represents the optimized protocol established after empirical testing of multiple incubation durations and temperatures. A schematic overview is provided in **Figure 1A**. Individual organoid systems may require minor adjustments (e.g., extended clearing for increased tissue size or density), but the workflow described represents a robust baseline for clearing of human brain organoids and assembloids.

**Figure 1.**
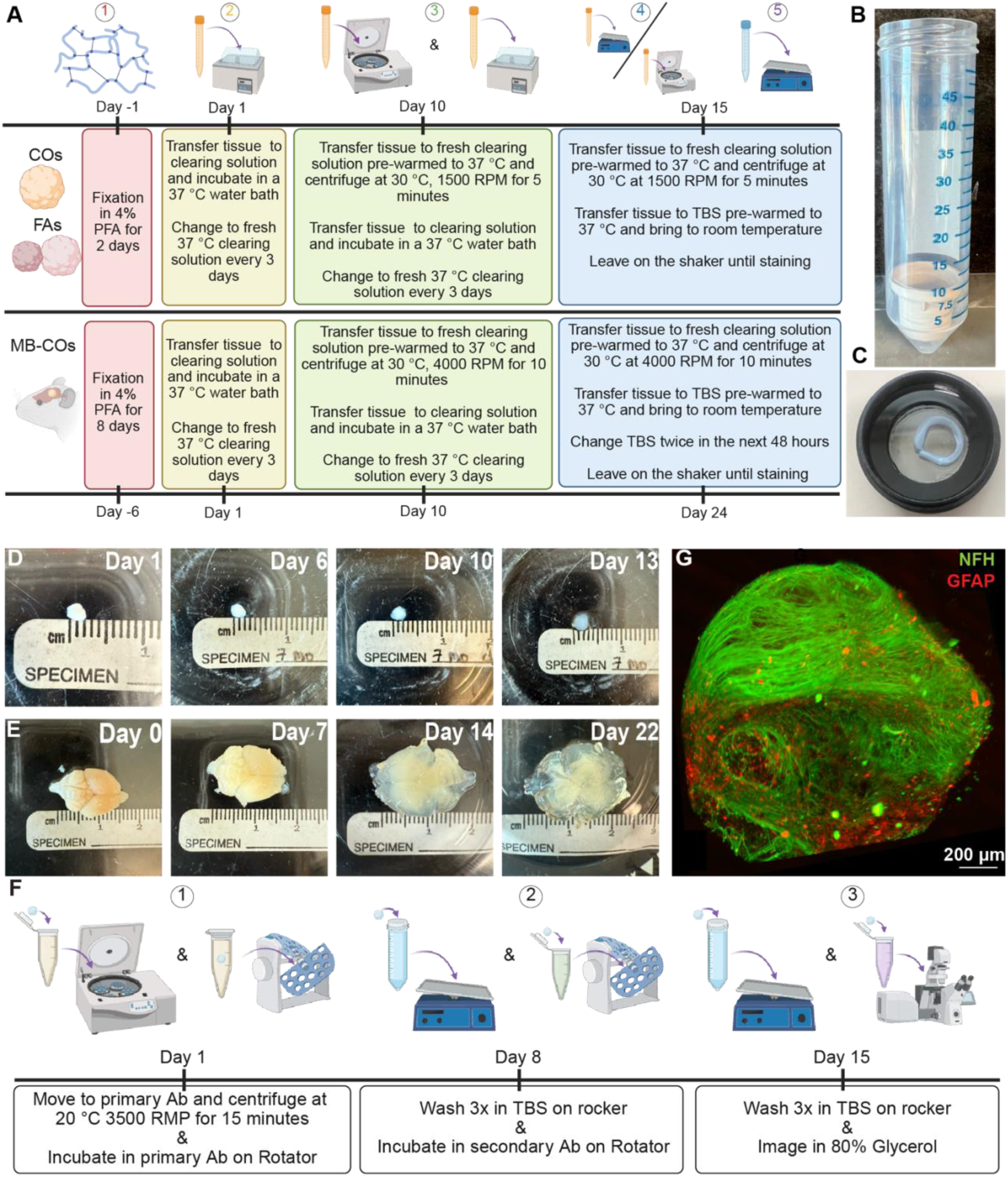
CLARI-O workflow for three-dimensional clearing and immunolabeling of human cortical organoids (COs), forebrain assembloids (FAs), and mouse brain-containing xenotransplanted cortical organoids (MB-COs). (A) Schematic overview of the CLARITY-O-clearing workflow optimized for COs, FAs, and MB-COs. COs, FAs, and MB-COs are first fixed in 4% paraformaldehyde, followed by incubation in clearing solution and sequential centrifugation steps to facilitate lipid removal and optical transparency. Samples are maintained at 37 °C with periodic replacement of clearing solution until sufficient tissue clearing is achieved prior to immunostaining. Illustration created with BioRender.com. (B) Representative image of Falcon tube fitted with a mesh insert. (C) Example of an assembly of the two glass-bottom PELCO wells, secured with BluTack to maintain stability and prevent dehydration throughout image acquisition. (D) Time-course images illustrating progressive optical clearing of CO during the CLARITY-O protocol. (E) Time-course images of MB-COs showing progressive tissue clearing over the course of the protocol. (F) Schematic of the downstream immunostaining workflow. Cleared samples are incubated with primary antibodies following centrifugation-assisted antibody penetration, washed in TBS, incubated with secondary antibodies, and subsequently prepared for imaging. Illustration created with BioRender.com. (G) Representative three-dimensional confocal reconstruction of a cleared day 120 CO immunolabeled for NFH (green; axonal marker) and GFAP (red; astrocytic marker), demonstrating preservation of neuronal and glial architecture following CLARITY-O processing. Scale bar, 200 µm.

### CLARI-O processing of mouse brains containing xenotransplanted cortical organoids (MB-COs)

Following transcardial perfusion, MB-COs were post-fixed in 4% paraformaldehyde for 8 days at 4 °C and then transferred to clearing solution in 50-mL Falcon tubes and incubated in a 37 °C water bath. The clearing solution was replaced every 3–5 days with freshly prepared, pre-warmed solution to ensure consistent detergent activity.

After 10 days of passive clearing, each MB-CO was transferred to a 50-mL Falcon tube fitted with a mesh Corning Netwell insert (cat. CLS3519, SigmaMillipore, **Figure 1B**), which suspends the tissue above the bottom of the tube, allowing uniform exposure to the clearing solution during centrifugation. Tubes were filled with 30 mL of pre-warmed clearing solution and centrifuged at 4000 rpm for 10 minutes. MB-COs were then returned to the 37 °C water bath and monitored daily. Once the desired level of optical transparency was achieved (usually at day 24), samples underwent a second centrifugation under the same conditions using a Falcon tube fitted with a mesh insert to remove any remaining detergent-rich solution.

Cleared brains were subsequently transferred into pre-warmed 0.05 M Tris-Buffered Saline (TBS) and washed at room temperature for 48 hours, with TBS replaced (always pre-warmed) at least twice during this period. Samples were then stored in 0.05 M TBS at room temperature on a shaker until immunostaining. A schematic overview of this protocol is also shown in **Figure 1A**.

#### Immunohistochemistry

Samples were placed in a 1.5 mL Eppendorf tube with primary antibody solution made with 0.05 M TBS, pH 7.6 and 0.5% Triton X-100 (cat. X100, MilliporeSigma). They were then centrifuged at 20 degrees Celsius for 15 minutes at 3500 rpm to increase antibody penetration. The following primary antibodies were used: rabbit polyclonal anti-GABA (1:500; Cat#A2052, Sigma Millipore), Chicken polyclonal anti-GFAP (1:1000; Cat# ab4674, Abcam, Waltham, MA, USA), Mouse polyclonal anti-GFAP conjugated to AlexaFluor-594 (1:1000; Cat#NBP1-05197AF594, Novus Biological, Littleton, CO, USA), Chicken polyclonal anti-Iba1 (1:500; Cat#IBA1, Aves Labs, Davis, CA, USA), Guinea pig polyclonal anti-MAP2 (1:500; Cat#188 004, Synaptic Systems, San Jose, CA, USA), Rabbit polyclonal anti-NeuN conjugated to AlexaFluor-594 (1:1000; Cat#NBP1-77686AF594, Novus Biological), Mouse monoclonal anti-NFH conjugated to AlexaFluor-488 (1:500; Cat#NB500-416AF488, Novus Biological), Mouse Monoclonal MBP Conjugated to ALexaFLuor 488 (1:500, Cat#NBP2-22121AF488, Novus Biological), Rabbit polyclonal anti-PSD95 (1:500; Cat#51-6900, Thermo Fisher Scientific), Goat polyclonal anti-PSD95 (1:500; Cat# ab12093, Abcam), Mouse monoclonal anti-Synaptophysin (1:1000; Cat#MAB368, Sigma Millipore).

Samples were incubated in 1.5-mL tubes for 7 days at room temperature. For primary antibodies that were unconjugated, the samples were washed in 50-mL conical tubes of 0.05 M TBS, pH 7.6 three times for 10 minutes each before being transferred to a secondary solution. The following secondary antibodies used: Donkey polyclonal anti-Mouse conjugated to AlexaFluor-488 (1:500; Cat#A-21202, Thermo Fisher Scientific), Goat polyclonal anti-Rabbit conjugated to AlexaFluor-568 (1:500; Cat#A-11011, Thermo Fisher Scientific), Donkey polyclonal anti-Rabbit conjugated to AlexaFluor-594 (1:500; Cat#A-21207, Thermo Fisher Scientific), Goat polyclonal anti-Guinea Pig conjugated to AlexaFluor-647 (1:500; Cat#A-21450, Thermo Fisher Scientific), Donkey polyclonal anti-Goat conjugated to AlexaFluor-647 (1:500; Cat#ab150135, Abcam). Antibodies at the indicated concentrations were added in Eppendorf tubes with 1.5 mL of solution consisting of 0.05 M TBS, pH 7.6 and 0.5% Triton X-100 for 7 days. If instead the samples were incubated with conjugated primary antibodies, this step was skipped. Finally, after unconjugated secondary/conjugated primary incubation, the samples were again washed three times for 10 minutes each in 0.05 M TBS, pH 7.6. They were then transferred to 80% glycerol (cat. EP229, ThermoFisher) in 0.05 M TBS, pH 7.6 for refractive index matching prior to imaging.

#### Lectin staining

DyLight 649 labeled tomato lectin (cat. DL11781, Vector Laboratories, Newark, CA, USA) was used to stain MB-CO for 48 hours following the manufactural instructions.

#### Imaging

Samples were mounted in 20% glycerol solution prepared in 0.05 M TBS (pH 7.6) and positioned between two glass-bottom PELCO wells, secured with Blu Tack to maintain stability and prevent dehydration throughout image acquisition (**Figure 1C**). High-resolution imaging was performed using a Leica TCS SP8 confocal microscope (Leica Microsystems, Wetzlar, Germany) equipped with a motorized XYZ stage and an HC FLUOTAR L 25× VISIR water-immersion objective (0.95 NA, 2.4 mm working distance), enabling deep optical sectioning of intact samples.

Detector settings and acquisition parameters were optimized individually for each sample to maximize signal-to-noise while preserving endogenous fluorescence. Image stacks were processed and rendered using either Leica LAS X software, or BITPLANE (IMARIS), including automated stitching for large fields of view and full 3D reconstruction of cleared organoids and tissue sections.

## Results

### CLARI-O enables rapid clearing and 3D imaging of intact COs and MB-COs

In 2019, Mortazavi et al.^17^ published a clearing method that not only allowed for the processing of whole rodent brains within 12 days but also proved being effective across multiple tissue types including human, non-human primate, and zebrafish brain. In this study, we further optimized Mortazavi’s approach by adapting it for use in COs, FAs, and MB-COs. A diagram of our organoid-optimized protocol, termed CLARI-O, is shown in **Figure 1A**, which provides a schematic timeline of this protocol, illustrating how the duration of each processing step is optimized according to whether it is a CO/FA, or MB-CO.

COs and FAs underwent fixation in 4% PFA for 2 days while MB-COs were fixed for 8 days. Following this, samples were incubated in 37°C clearing solution for 10 days with fresh pre-warmed solution changes every 3 days. We prioritized passive clearing over electrophoretic tissue clearing (ETC) as the passive method is milder and preserves the integrity of small, fragile tissue samples^17^. A key modification we introduced to the 2019 Mortazavi et al. protocol is the inclusion of a centrifugation step after the 10-day incubation phase for all samples. This centrifugation step significantly enhances the infiltration of the clearing solution into the tissue and leads to the removal of residual detergent-rich solution that can prevent uniform transparency leading to clearing related artifacts. After centrifugation, COs and FAs were returned to 37 °C clearing solution for 5 days while MB-COs were returned for 14 days. In both cases, the solution was again changed with pre-warmed solution every 3 days. On the final day of heat-assisted clearing (Day 15 for COs and FAs and Day 24 for MB-COs) the samples were centrifuged again and then transferred to pre-warmed TBS, brought to room temperature and left of the shaker until staining.

Using this protocol, we achieved optical transparency in COs and FAs in under 15 days, and in MB-COs in under 24 days (**Fig. 1D and 1E**). Following clearing, CO and FA samples were incubated in primary and secondary antibody solutions for one week each. Like the clearing phase, we utilized centrifugation at the start of the primary incubation to facilitate antibody penetration. **Figure 1F** details the immunohistochemistry timeline for COs and FAs. Following clearing and staining, we use 2 PELCO glass bottom dishes and Blue Tak (**Figure 1C**) for confocal imaging. To do this we first prepared a well with Blue Tak on one dish. The tissue is placed within this well, and a solution of 80% glycerol is added. The second Pelco dish then sits on top of this well. A circular bubble level is used to make sure the tissue sandwich is leveled prior to imaging. This also allows for both sides and orientation of the glass dishes to be imaged (**Figure 1C**). Furthermore, this process allowed us to maintain sample stability and image the entire organoid by forgoing the need for classical histological sectioning which inevitably severs neuronal processes. Capitalizing on this preservation of tissue integrity, we successfully visualized a whole cleared day-120 CO stained for Neurofilament-heavy chain axonal marker (NF-H) and the astrocytic marker GFAP, revealing the directionality and spatial distribution of neuronal axons and astrocytes within the intact 3D architecture (**Figure. 1G** and **Movie file 1**).

### Visualization of cellular heterogeneity in COs using CLARI-O

One of the caveats of classical CO models is a limited heterogeneity and the lack of specific glia cells, such as oligodendrocytes and microglia. Recent advancements allow us to overcome these limitations^12,21^. By introducing PDGF-AA, IGF, and T3, we successfully expanded the glial progenitor pool and directed their differentiation into oligodendrocyte progenitor cells (OPCs) and mature oligodendrocytes within dorsally-derived oligodendrocyte-containing COs (dOCOs)^12^. We applied CLARI-O to clear day 120 dOCOs and immunostained them for NeuN, neuronal soma marker, microtubule-associated protein 2 (MAP2), labeling dendrites, and myelin basic protein (MBP), a marker of mature oligodendrocytes. Confocal imaging and 3D reconstruction revealed that oligodendrocytes were distributed in discrete, patchy clusters throughout the dOCO (**Figure. 2A, B**). This distribution likely reflects the clonal expansion of distinct glial progenitor niches that cannot be clearly observed in MBP-stained dOCO sections.

**Figure 2.**
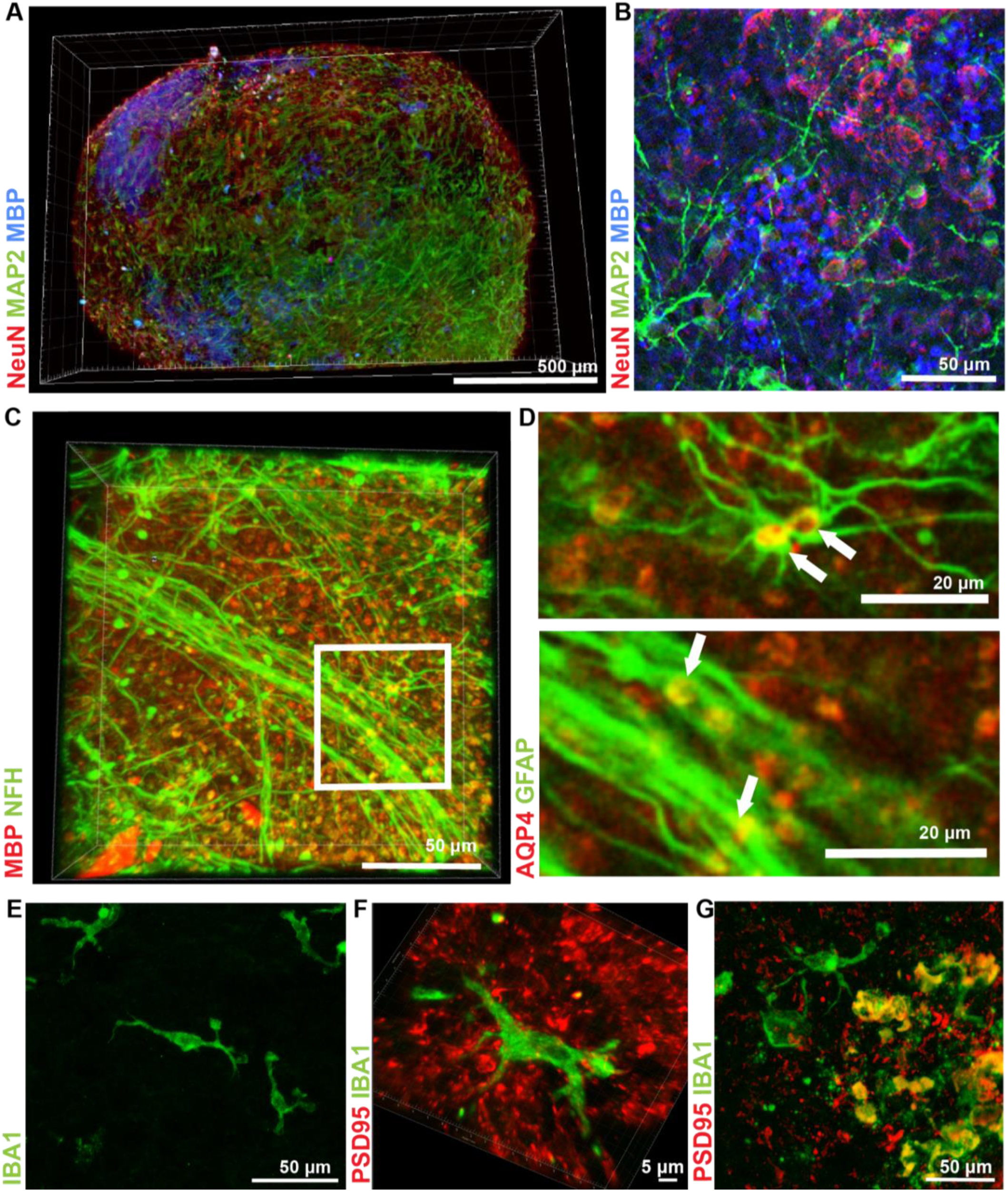
CLARI-O three-dimensional imaging of human COs reveals neuronal, myeline basic protein positive oligodendroglial, astrocytic, and microglial organization. (A) Volumetric reconstruction of a CLARITY-O day 150 OCO immunolabeled for MAP2 (green; dendritic, neuronal marker), MBP (blue; myelin basic protein), and neurons (red). The 3D rendering demonstrates extensive neuronal architecture with interspersed MBP-positive oligodendroglial niches throughout the organoid volume. Scale bar, 500 µm. (B) Higher-magnification view with inset highlighting a localized pool of MBP-positive oligodendrocytes within the parenchyma. Scale bar, 50 µm. (C) Low-magnification view of co-labeling for NFH (axonal marker) and MBP demonstrates MBP-positive oligodendrocytes in close association with NFH-positive axonal fibers (white arrows), consistent with axon–glial interactions within the CO. Scale bar, 50 µm. The boxed region indicates the area shown at higher magnification in the insets in D. (D) Upper inset: Higher-magnification view demonstrating an MBP-positive oligodendrocytes (arrows) that associate with nearby NFH-positive axons. Individual axons can be resolved as they become ensheathed by oligodendrocyte processes, consistent with active myelination. Lower inset: High-magnification view of a parallel axonal bundle composed of NFH-positive axons (green), with MBP-positive oligodendrocytes positioned within the tract (arrows). (E) IBA1-positive microglia (green) incorporated into day 120 CO 7 days prior to fixation. Scale bar, 50 µm. (F, G) IBA1-positive microglia (green) and PSD95 (red) labeling demonstrate microglial proximity to synaptic structures, suggesting active microglial–synaptic interactions with synaptic material and a functional immune surveillance state. Scale bar, 5 µm in F and 50 µm in F.

Furthermore, to investigate axon-glia interactions, we co-stained for NF-H. 3D reconstruction demonstrated colocalization between MBP+ oligodendrocytes and NF-H+ axons, indicating points of oligo-axonal contact and potential ensheathment or myelination (**Figure. 2C**, white arrows). Higher-magnification views (**Figure. 2D**) reveal oligodendrocyte processes directly ensheathing individual axons, as well as MBP-positive cells positioned within organized axonal bundles, consistent with active, coordinated myelination.

Conventional 2D histological approaches are limited in their ability to capture the spatial extent required to follow long axonal processes and the full scope of oligodendrocyte interactions along them. In contrast, CLARI-O preserves the intact 3D architecture, enabling visualization of axon–glia relationships across extended volumes. Notably, even though we detected MBP-positive NF-H colocalized oligodendrocytes, this spatial overlap was relatively modest (**Figure. 2C, D**). Given that microglia originate from the mesoderm, which is distinct from the neuroectodermal lineage of cortical organoids, they are not typically found in COs^25^. Recent studies showed that the introduction of microglia into organoids shaped the neuronal activity of the organoids, and emphasized the critical role of microglia in shaping neuronal activity through synaptic surveillance and refinement^23,26^. Considering their crucial role, we separately differentiated iPSC into microglia and subsequently incorporated iPSC-derived microglia into COs^24,27^. COs were fixed 7 days following the incorporation and processed using CLARI-O. Immunohistochemical staining for microglia (IBA1) followed by confocal 3D reconstruction, demonstrated the effective infiltration of microglia into the CO (**Figure 2E-G** and **Movie file 2**). Crucially, we observed a significant colocalization between IBA1+ microglial processes and PSD-95+ synaptic puncta, indicating that these microglia actively interact with the synaptic material (**Figure 2 F, G**). Importantly, conventional sectioning often obscures full microglial morphology, frequently severing fine processes depending on the cutting plane. This limitation is especially pronounced in COs, where microglia are sparse, unevenly distributed, and their locations are unknown prior to sectioning, making intact capture challenging. In contrast, CLARI-O as a section-free approach enables visualization of multiple intact microglia with diverse morphologies (**Figure 2E-G**) and allows for the visualization of their physical interaction with synaptic elements in a 3D environment.

### Section-Free Visualization of Forebrain Assembloid Fusion and Interneuron Integration Using CLARI-O

Beyond single COs, we applied CLARI-O to visualize the complex architecture of forebrain assembloids (FAs) generated by the fusion of ventral (vCO) and dorsal (dCO) organoids. While the migration of GABAergic interneurons and oligodendrocytes across these boundaries is well-documented, the specific structural mechanisms facilitating this fusion and migration remains poorly understood.

Immunostaining of CLARI-O-processed FAs for NeuN and GFAP revealed, for the first time, the existence of GFAP+ parallel fibers specifically located at the junction between the dCO and vCO (**Figure 3A, B**). These fibers appear to serve as a scaffold, providing structural support for migrating NeuN+ neurons (white arrows in **Figure 3B**). The presence of these glial bridges at the interface of two tissues is highly reminiscent of glial scar formation observed following CNS injury^28^. To our knowledge, this is the first detailed characterization of the structural components—specifically GFAP+ fibers, which may be required for physical assembloid fusion.

**Figure 3.**
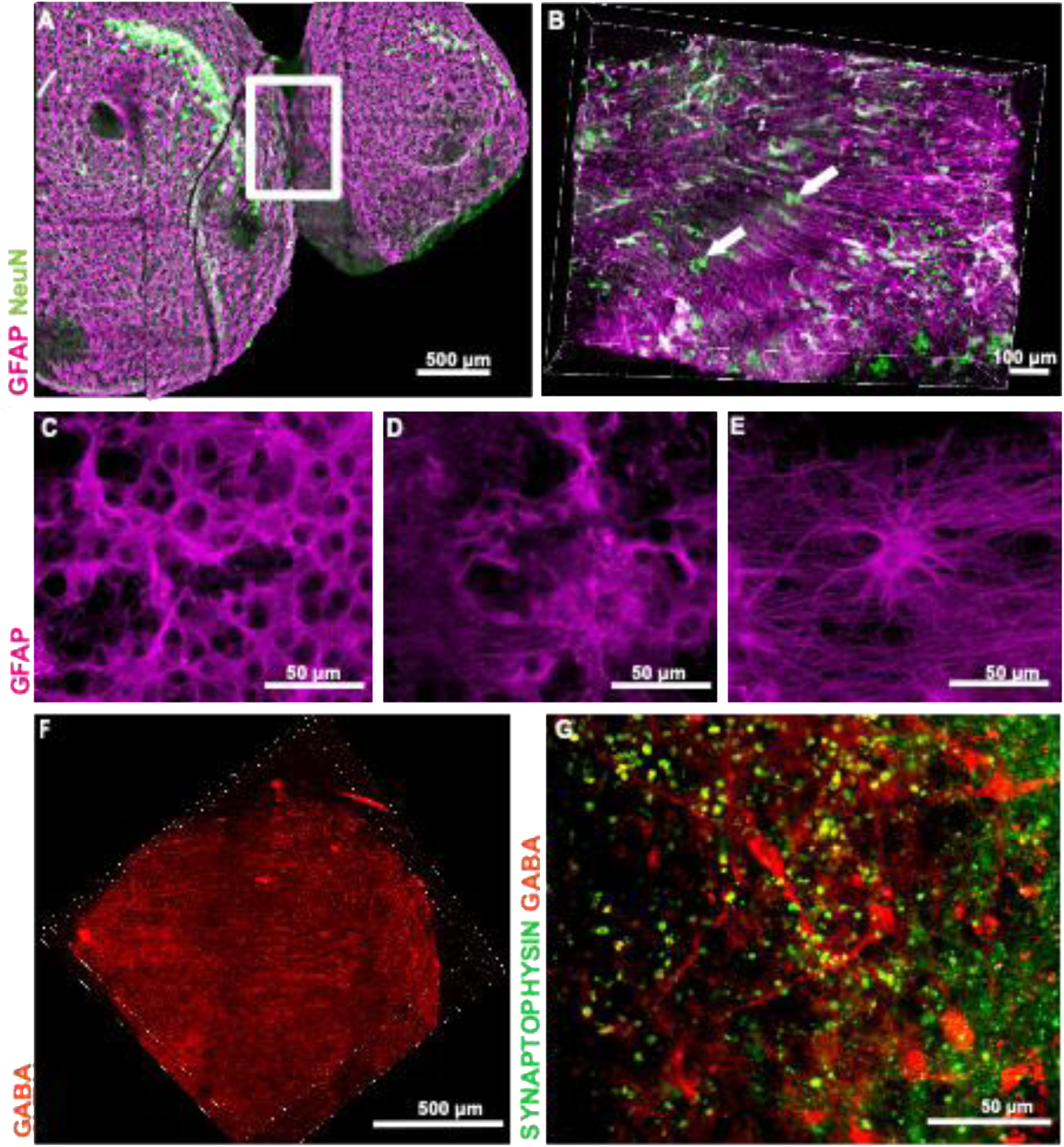
Three-dimensional CLARI-O imaging of human forebrain assembloids (FAs) reveals neuronal migration, astrocytic heterogeneity, and GABAergic network integration. (A) Low-magnification view of a CLARI-O-cleared day 120 FA immunolabeled for NeuN (green; mature neurons) and GFAP (magenta; astrocytes). The composite image demonstrates large-scale neuronal distribution embedded within an extensive astroglial scaffold. Scale bar, 500 µm. (B) Higher-magnification rendering of the boxed region in (A), showing NeuN-positive neurons (white arrows) interspersed within and migrating along GFAP-positive astrocytic fibers, consistent with astroglia-guided neuronal migration within the FAs. Scale bar, 100 µm. (C–E) Representative images illustrating three distinct astrocytic morphologies observed within the assembloid: (C) densely packed, stellate astrocytes with overlapping processes; (D) transitional astrocytes with elongated somata and moderate branching; and (E) highly ramified astrocytes with long, radially oriented processes forming a fibrous network. Scale bars, 50 µm. (F) Three-dimensional rendering of GABA-positive interneurons and their neurites within the assembloid, demonstrating extensive process outgrowth and migration across the tissue volume. Scale bar, 50 µm. (G) High-magnification image of GABA-positive cells (red) exhibiting developed neuronal morphology and complex arborization. Synaptophysin (green) labels presynaptic puncta; regions of overlap suggest synaptic vesicle compartments potentially containing GABA, consistent with functional maturation of inhibitory circuits. Scale bar, 50 µm.

In addition, we also visualized a morphologically distinct population of astrocytes within FAs. As shown in **Figure 3C-E**, astrocytes exhibit marked heterogeneity in soma size, process length, and branching architecture, reflecting the emergence of territorial organization and ongoing astrocytic maturation^29^. We identified dense mesh-like networks of closely packed astrocytes with thick, highly branched processes and substantial overlap, consistent with immature or early protoplasmic-like states (**Figure 3C**); transitional astrocytes with enlarged somata, thicker processes, and partial radial organization suggestive of intermediate maturation (**Figure 3D**); and stellate astrocytes characterized by well-defined somata and long, thin, radially oriented processes extending across broader domains, indicative of advanced structural complexity (**Figure 3E**)^30,31^.

FAs serve as an ideal model for studying inhibitory GABAergic interneurons, the majority of which originate in the ventral telencephalon and migrate dorsally to populate the cortex, hippocampus, and amygdala^32,33^. These interneurons are critical for cortical development, as they refine neuronal circuitry and modulate spontaneous synchronous activity^34,35^. Using CLARI-O together with immunohistochemistry, we visualized GABAergic interneurons integrated within the surrounding synaptic circuitry, where synaptophysin (SYN⁺) vesicles were frequently associated with GABA⁺ neurites and somata, suggesting functional incorporation of inhibitory neurons into the developing network (**Figure 3H, G**).

### Whole-brain clearing enables 3D visualization of xenograft maturation and integration within the mouse cortex

Absence of vasculature and limited oxygen and nutrient delivery as well as restricted neuronal maturation are some of the limitations of organoid models *in vitro*. To address these challenges, we and others have developed and optimized xenotransplantation of COs into the rodent brain^9,10^. When engrafted into the mouse retrosplenial cortex (**Figure 4A, Ai**), COs become vascularized by host vasculature, achieve structural and functional integration, and display robust morphological and functional maturation^9^. Clearing xenotransplanted COs within the intact mouse brain is therefore critical for obtaining a comprehensive view of their spatial organization, holistic visualization of cell migration away from the implantation site, and vascular integration.

**Figure 4.**
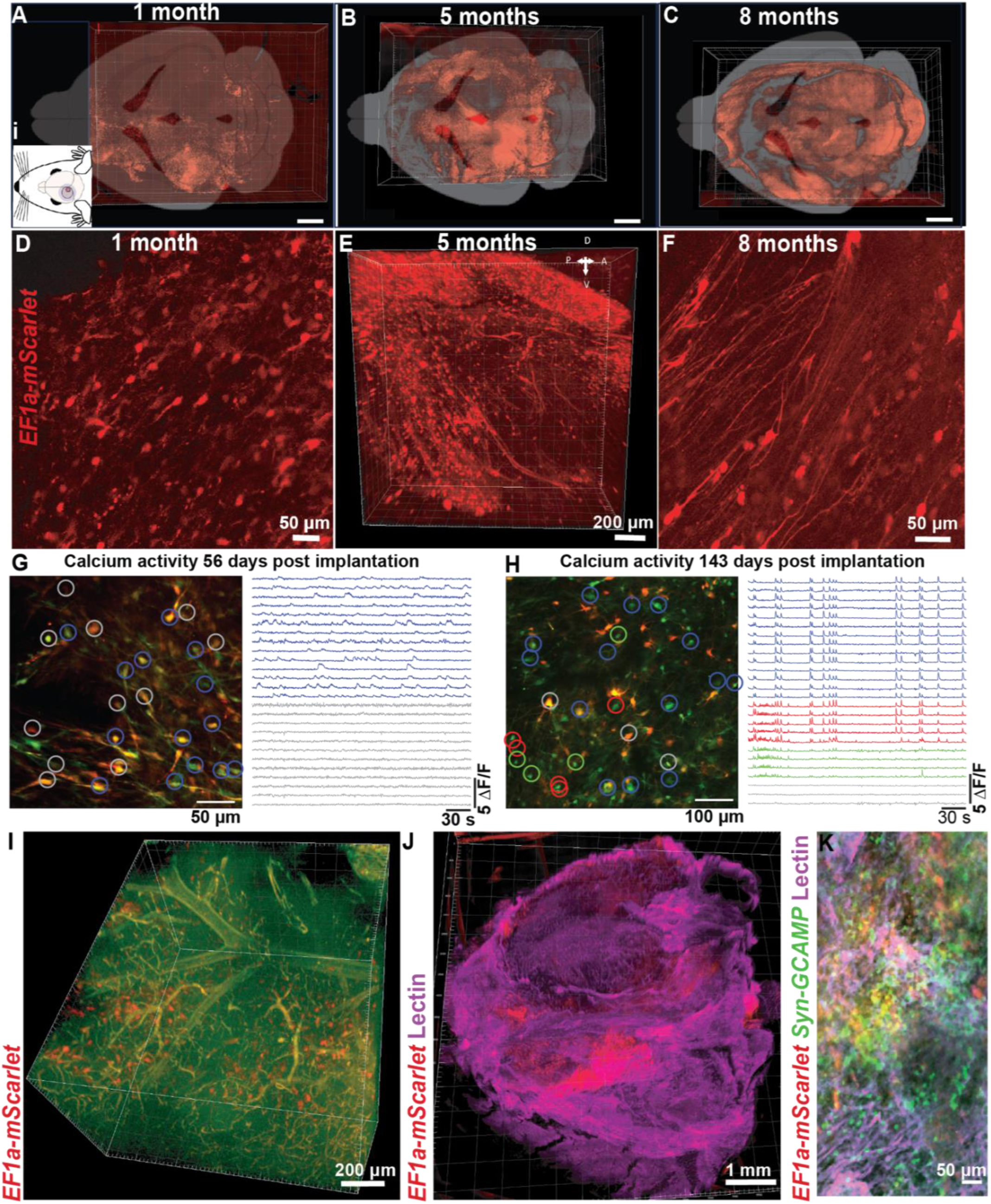
Long-term survival, integration, functional maturation, and vascularization of MB-COs visualized with CLARI-O. The composite image demonstrates (A–C) Volumetric three-dimensional reconstructions of MB-COs virally labeled with *EF1a-mScarlet* (red) showing the spatial distribution of cells within the host brain at 1 month (A), 5 months (B), and 8 months (C) after implantation. Renderings from different post-transplantation time points illustrate progressive growth, structural integration, and spatial distribution of graft-derived cells within the host tissue. Scale bars, 1 mm. (D) Higher-magnification view of the graft region at an early post-transplantation stage showing mScarlet-positive human cells concentrated near the graft core with relatively simple cellular morphologies characterized by short neurites and limited branching. Scale bar, 50 µm. (E–F) Higher-magnification views at later post-transplantation stages showing increased dispersion of graft-derived cells throughout the surrounding host tissue. Cells exhibit pronounced morphological maturation with elongated neurites, increased branching complexity, and more elaborate neuronal arborization. Scale bar, 200 µm (E) and 50 µm. (G, H) In vivo calcium imaging of spontaneous neuronal activity using GCAMP8s-labeles human CO-derived neurons 56 (G) and 143 (H) days after transplantation. Left panels show field of view with individual neurons outlined; representative ΔF/F calcium traces are shown on the right, Scale bar, 50 µm (G) and 100 µm (H). Calcium imaging at 143 days post implantation demonstrating increased neuronal activity and more synchronized network dynamics among transplanted ne5urons (H) as compared to the earlier time point (G). (I) Three-dimensional reconstruction of the graft region demonstrating extensive host vascularization surrounding and penetrating the transplanted tissue, visualized through intrinsic vascular autofluorescence. Scale bar, 200 µm. (J) Whole-graft volumetric reconstruction demonstrating vascularization of the transplanted CO using lectin labeling (magenta) of host vasculature. Human graft cells are labeled with EFa1a-mScarlet. Scale bar, 1 mm. (K) High-magnification image showing transplanted human CO-derived neurons labeled with *Syn-GCaMP* (green) and *mScarlet* (red) and vasculature labeled with lectin (magenta). Scale bar, 50 µm.

Here, we demonstrate that intact whole mouse brains can be safely and reproducibly processed using CLARI-O, following the workflow outlined in **Fig. 1A**. One of the key advantages of CLARI-O is its compatibility with endogenous fluorescent reporters and live-cell imaging workflows. Xenotransplanted COs labeled prior to transplantation with mScarlet under a ubiquitous promoter via lentiviral transduction (LV-EF1alpha-mScarlet) retained robust fluorescent signals at 1, 5, and 8 months post-xenotransplantation (**Fig. 4A-F**). This signal remained readily detectable by confocal microscopy following CLARI-O processing.

Importantly, CLARI-O enables whole-brain visualization of how xenotransplanted CO cells structurally integrate and disperse within the host brain over time. By combining CO xenotransplantation with whole-brain CLARI-O clearing and volumetric imaging, we were able to directly track the spatial distribution and morphology of human CO-derived cells throughout the intact mouse brain. At 1-month post-transplantation, mScarlet-positive human cells were largely confined to the implantation site and exhibited relatively immature morphologies, characterized by short neurites and limited branching (**Figure. 4A, D** and **Movie files 3, 4**). In contrast, by 5 months post-transplantation, CO-derived cells were markedly more dispersed, migrating well beyond the graft core and in some cases reaching distant regions, including the contralateral hemisphere **(Figure. 4B, E, F)** and cerebellum at 8 months (**Figure. 4C**). These later stages were accompanied by pronounced morphological maturation, with transplanted cells displaying elongated neurites, increased branching complexity, and elaborate arborization patterns consistent with advanced neuronal differentiation and integration into host circuitry **(Fig. 4E, F).** Together, these data demonstrate that whole-brain CLARI-O imaging provides a powerful platform to map the large-scale migration and maturation of transplanted human neurons within the intact host brain.

To examine the relationship between structural maturation and functional integration following xenotransplantation, we performed longitudinal assessments of neuronal activity using calcium imaging with the GCaMP8s sensor *in-vivo* prior to perfusion. Like the mScarlet labeling strategy, COs were transduced with lentivirus to drive neuron-specific expression of GCaMP8s under the synapsin promoter (LV-hSyn1-GCaMP8s). Consistent with our prior work, this approach enables longitudinal monitoring of calcium dynamics in transplanted neurons for up to 8 months post-transplantation, using a range of in vivo imaging modalities^9,36^.

Using two-photon (2P) microscopy, we quantified fluorescence changes in neuronal somata and extracted single-cell activity traces, which were expressed as ΔF/F^36^. As shown in **Figure 4G**, at 56 days (∼1 month) post-transplantation, GCaMP8s-labeled neurons exhibited largely asynchronous and disorganized calcium transients. In contrast, as seen in **Figure 4H**, by 143 days (∼ 5 months) post-transplantation, neuronal activity displayed a marked increase in synchrony across most recorded neurons, indicative of progressive functional maturation and network-level integration^37^. These functional measurements complement spatial and structural changes observed through CLARI-O in human transplanted CO-derived neurons detected with mouse cortex.

Finally, we used CLARI-O processed MB-COs to visualize the extent of graft vascularization within the mouse cortex. Vascularization was assessed using an autofluorescence-based approach previously validated in zebrafish^17,38^ (**Figure. 4I**), as well as by lectin staining performed on MB-COs at 8 months post-transplantation (**Figure. 4J, K**). Lectin binds to sugar residues present on endothelial cells and serves as a marker of vascular structures^39^. Confocal imaging revealed extensive vascularization of the graft by the host mouse vasculature. **Figure 4K** shows human, mScarlet and GCAMP-labeled cells embedded within the mouse vasculature (stained with lectin). Collectively, we demonstrate the strength of CLARI-O as a platform for resolving graft vascularization and human cell integration within the mouse cortex in an intact three-dimensional context.

## Discussion

Traditional 2D cell cultures lack the spatial and structural complexities present in living tissues, making them insufficient for studying realistic cellular interactions. In 3D models, cells preserve the spatial cues and cell-matrix interactions necessary to maintain their natural structures, polarities, and behaviors. Human iPSC-derived COs have been invaluable in modeling brain structures and cellular interactions in healthy and disease-associated brain. However, despite these advances, structural and phenotypic analyses following functional assessments, still rely on slicing fixed, frozen COs for subsequent immunostaining. This approach undermines the benefits of the 3D architecture by reducing the data into 2D slices, disrupting spatial orientation, and severing axonal and dendritic projections.

In the present study, we introduce CLARI-O, an optimized clearing workflow specifically adapted for COs, forebrain assembloids (FAs), and mouse brains containing xenotransplanted organoids (MB-COs). CLARI-O incorporates low-speed centrifugation and sustained heat-assisted passive clearing, a combination that, to our knowledge, has not been systematically applied to human brain organoids or assembloids. This approach meaningfully accelerates lipid extraction while minimizing mechanical disruption, enabling highly consistent clearing across samples of varying size and developmental complexity. The brief centrifugation step helps remove residual detergent-rich solution trapped within the samples, thereby improving the uniformity of transparency and reducing clearing artifacts that commonly arise in spherical organoids. Maintaining all clearing solutions at 37 °C prevents fluctuations in lipid solubilization kinetics^17^, which is particularly important for organoids whose dense cytoarchitecture can hinder passive diffusion. This centrifugation-enhanced, heat-stabilized passive clearing protocol standardizes and streamlines the clearing process while preserving delicate 3D architecture.

Using post-clarity immunostaining and high-resolution confocal imaging, we provide a step-by-step approach to visualize specific cellular structures as well as neuronal and glial cell types. Furthermore, we extended our protocol to MB-COs to enable the visualization of the integration of human cells into the host brain at different stages and extensive vascularization of the xenotransplanted COs within the mouse brain.

We further applied CLARI-O to dOCOs containing oligodendrocytes generated using our previously published differentiation protocol. This protocol targets a small endogenous population of oligodendrocyte progenitor cells and expands their pool over time. Visualization of CLARI-O-processed and immunolabeled dOCOs revealed a patchy distribution of MBP-positive oligodendrocytes, consistent with their derivation from a limited number of glial progenitors. Although MBP-positive oligodendrocytes were readily detectable in day 150 dOCOs, their colocalization with axonal markers is relatively limited, suggesting a potentially constrained myelination capacity within this type of model and a more prolonged organoid maturation that may be necessary to achieve more robust and extensive myelination. Imaging of incorporated microglia enabled high-resolution visualization of their morphology and revealed microglia–synapse interactions, supporting their role in synaptic refinement.

Assembloid systems have emerged as powerful tools for modeling interregional connectivity in the developing brain^40,41^. Previous studies showed multimodal migration of ventral organoid–derived interneurons across fusion interfaces^42^ and it has been proposed that the alignment of the progenitor zones at the interface of the fused organoids provides a permissive migratory environment^22,42^. Using CLARI-O-processed FAs, we visualized, for the first time, continuous radial glia GFAP-positive scaffolds spanning the fusion interface. In conjunction with previous works that show axon-level integration and migration of oligodendrocytes and interneurons through the fusion midline^22,43,44^, our findings provide structural evidence for a glial substrate that may facilitate inter-organoid cellular migration and integration. We also demonstrated a morphologically heterogenous population of astrocytes spanning different developmental stages.

A major advance of this study is the successful clearing of entire mouse brains containing fluorescently labeled transplanted human organoids. We visualized the integration of the human cells into mouse tissue at 1, 5, and 8 months post-surgery. Following the established pipeline to monitor neuronal activity over time using a GCAMP8s calcium sensor, we showed the increased synchrony of neuronal firing over time. In intact human brain, this synchrony picks up before birth and is critical for the proper development of neuronal networks and enhances activity-dependent synaptic refinement^45–47^. This shift in spontaneous neuronal activity was accompanied by increased morphological complexity of human neurons, as well as their dispersion and migration within mouse cortex. Moreover, CLARI-O-based imaging revealed a substantial expansion of fluorescently labeled human neurons over time, with some cells potentially acquiring callosal identities and projecting to the contralateral hemisphere. These structural findings should be complemented in future studies by immunohistochemical staining to refine cell-type identification. Finally, lectin labeling further demonstrated robust vascularization of the xenotransplanted COs, consistent with prior observations of extensive vascular integration in vivo^9,10,36^. The dense vascular network of host-derived blood vessels is likely essential for long-term graft survival, integration, and maturation, since it supports the supply of oxygen and nutrients, promotes metabolic exchange, and facilitates the maturation of CO-derived neurons.

To our knowledge, only few studies applied clearing protocol to assess phenotypic and structural properties of brain organoids, and none has presented a protocol suitable for clearing of MB-COs. Two studies utilized CLARITY, an acrylamide-based hydrogel embedding approach in combination with light-sheet microscopy^14,48^. Sakaguchi et al, 2019^48^ focused on calcium transients in cerebral organoids and used light sheet miscopy and CLARITY clearing to show the presence of VZ-like structures. However, the study provided limited methodological detail regarding the clearing procedure itself. Shnaider & Pristyazhnyuk., 2021^14^, on the other hand, described a more detailed protocol for CLARITY-based processing, but focused on a single stain of deep layer neurons that required agarose embedding optimized for light sheet microscopy. Most recently, Lange et al. (2025)^18^ used microglia-integrated brain organoids to model myelin repair and applied ScaleS4^49^, a hydrophilic clearing method that results in a comparatively limited lipid removal. A major drawback of using ScaleS4 is reduced tissue transparency relative to methods that achieve more extensive lipid removal^49^.

To overcome these limitations, we adapted an acrylamide-free passive clearing strategy originally developed by Xu et al. 2018^50^ and optimized by Mortazavi et al., 2019 to enhance tissue permeabilization^17^. Several features make this approach particularly suitable for COs, FAs, and MB-COs. Omitting acrylamide embedding increases antibody penetration^51,52^, does not result in protein loss as compared to acrylamide embedding^53^ and reduces tissue expansion^53^. Furthermore, omission of acrylamide embedding leads to a more streamlined protocol that reduces the processing time while preserving the native tissue structure^25,51^. Our “passive” form of clearing also omits electrophoresis (often used in other clearing approaches), utilizing passive immersion, that requires no specialized electrophoretic equipment and reduces the risk of tissue damage from heat or bubbles. As compared to the Mortazavi et al., 2019 protocol^17^, CLARI-O integrates low-speed centrifugation and extended heat-assisted passive clearing into the workflow to achieve uniform transparency, while preserving the complex and delicate architecture of COs and FAs as well as of both host and CO graft tissue in MB-COs.

Although we focused here on CLARI-O-processed FAs, our described methods can be straightforwardly transferred to other assembloid models such as corticothalamic assembloids, corticostriatal assembloids, and cortical-spinal-muscle assembloids^54^. We also demonstrated for the first time CALRI-O based clearing and staining of xenotransplanted organoids following functional in-vivo analysis using multiphoton imaging. In the future, we will explore the use of CLARI-O to correlate the identity of cells with certain functional behaviors, i.e., calcium activity patterns, with post-mortem structural analyses such as in-situ immunostaining using endogenous fluorescence or vascular structures as landmarks to co-register *in vivo* and post-mortem imaging datasets. This protocol is also compatible with downstream quantitative analyses and can complement multielectrode array recordings, optogenetic stimulation, and other functional assays.

In summary, our CLARI-O provides a robust, scalable, and accessible platform for high-resolution three-dimensional visualization of organoids, assembloids, and xenografts, bridging the gap between structure and activity and providing an invaluable resource for exploring disease mechanisms, testing drug effects, and developing new therapeutic strategies in next-generation 3D neural models.

## Limitations of the study

In the current study we did not perform detailed structural or morphometric quantification of the cleared COs and FAs. This limitation is partly related to the relatively small sample size in this study, particularly for MB-CO) samples, which makes quantitative comparisons across time points derived from individual samples or animals challenging. Nevertheless, these clearing and imaging techniques *can* support such analyses, offering a more efficient and accurate approach to 3D quantification that avoids the limitations of traditional 2D histological methods. This method can also be correlated with functional assessments of human neurons to provide insight into their progressive maturation within the neurodevelopmentally supportive environment of the mouse brain^9,36^.

In addition, our current implementation is relatively low throughput, as we performed a single round of immunostaining per organoid, whether derived *in vitro* or recovered following xenotransplantation. However, this reflects an experimental choice rather than a technical limitation of the CLARI-O workflow, and iterative staining and re-staining^55^ should be feasible with this approach. The primary constraint in our study was the limited number of fluorescent channels available on our imaging system (three), which restricted multiplexed marker detection. Access to microscopy platforms with expanded spectral capacity would substantially enhance the depth and multiplexing potential of CLARI-O–based analyses.

A further practical limitation of volumetric CLARI-O imaging is the substantial imaging and computational infrastructure required to acquire, process, and visualize large-scale 3D datasets from cleared tissues. Because intact COs and MB-COs are imaged without sectioning, optical access through large tissue volumes requires the use of long-working-distance objectives that are compatible with refractive-index–matched clearing media. In addition, whole-sample imaging generates very large datasets due to tiled acquisition across the cleared specimen. For example, a single organoid imaged at 20X magnification typically required ∼60 tiled scans (1024 × 1024 pixels each), with acquisition times approaching 10 hours and datasets of approximately 60 GB. Imaging of MB-COs produced even larger datasets due to the increased tissue volume and higher magnification required to resolve cellular structures. Whole-brain MB-CO scans at 10× magnification required 324 tiled images (512 × 512 pixels, triple-channel acquisition), resulting in approximately 13 hours of imaging and datasets exceeding 68 GB per sample. Processing, stitching, and rendering image volumes of this scale required a custom-built high-performance workstation with expanded RAM, high-capacity storage, and dedicated GPU resources to enable efficient 3D visualization. Although these requirements currently limit throughput and accessibility, rapid advances in imaging hardware, data storage, and GPU-based visualization platforms are expected to substantially improve the scalability of CLARI-O–based volumetric analyses.

## Conflict of Interest Statement

All authors declare that they have no conflicts of interest.

## Acknowledgments

We are grateful to NIH agencies for their support: NIH/NIA RF1AG088529(MPI: E. Zeldich and M. Thunemann), NIH/NINDS R21NS125469 (MPIs: E Zeldich and M. Medalla), NIH/NIA R21AG080269 (PI: E. Zeldich); NIH/NIA: R03NS126864 (PI: E. Zeldich). SB was supported by Medical Student Summer Research Program, Boston University. Figures were created with BioRender.com

We thank members of the Devor/Thunemann lab for technical support, Dr. Omer Revah (Hebrew University of Jerusalem) for his invaluable advice regarding viral labelling strategies, Dr. Steven J. Haggarty (Harvard Medical School and Massachusetts General Hospital) for kindly providing the MGH2046 hIPSC line, and the Boston University Neurophotonics Center for access to imaging facilities and technical support.

## Declaration of generative AI and AI-assisted technologies in the writing process

During the preparation of this work the authors used in ChatGPT (OpenAI) to assist with language editing, proofreading, and improving clarity of the manuscript. After using this tool, the authors reviewed and edited the content as needed and take full responsibility for the content of the published article.

## Author contributions

SB, FM, and EZ designed and performed experiments, gather data, wrote and edited the manuscript. FM and EZ conceived the idea, designed experiments, provided guidance, and supervised. SB, NBC, EK, and SKA designed and performed experiments related to the generation of organoids, incorporation of glia cells, staining and imaging. MT led experiments and analysis related to the xenotransplantation of the organoids. All authors read and edited the final manuscript.

## Notes

### Competing Interest Statement

The authors have declared no competing interest.

